## Supplementary material. Figures S1-S6 and Tables S1-S5. for "Comprehensive profiling of polyclonal sera targeting a non-enveloped viral capsid"

Figures S1-S6 and Tables S1-S5.

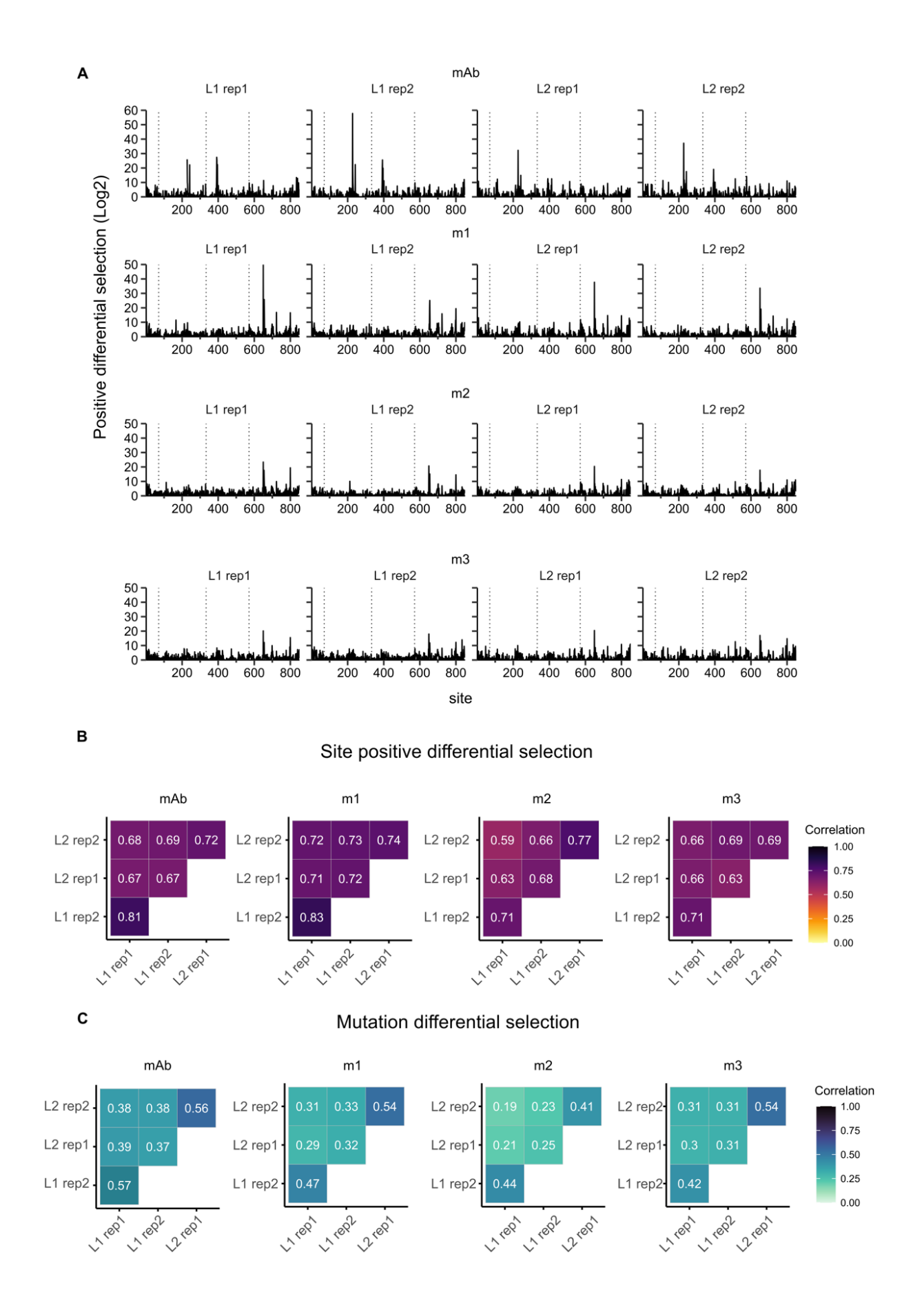

**Figure S1. Related to Figures 1 and 2. Technical and biological replicates of the mutational antigenic profiling of a mAb and mouse sera. (A)** Positive differential selection profiles of antigenic profiling of the mAb and three mouse sera samples (m1, m2, and m3). Two replicates (rep1 and rep2) were performed for two independent virus libraries: library 1 and library 2 (L1 and L2, respectively). **(B-C)** Matrix of Pearson correlation coefficients of site positive differential selection **(B)** and mutation differential selection **(C)** between biological and technical replicates.

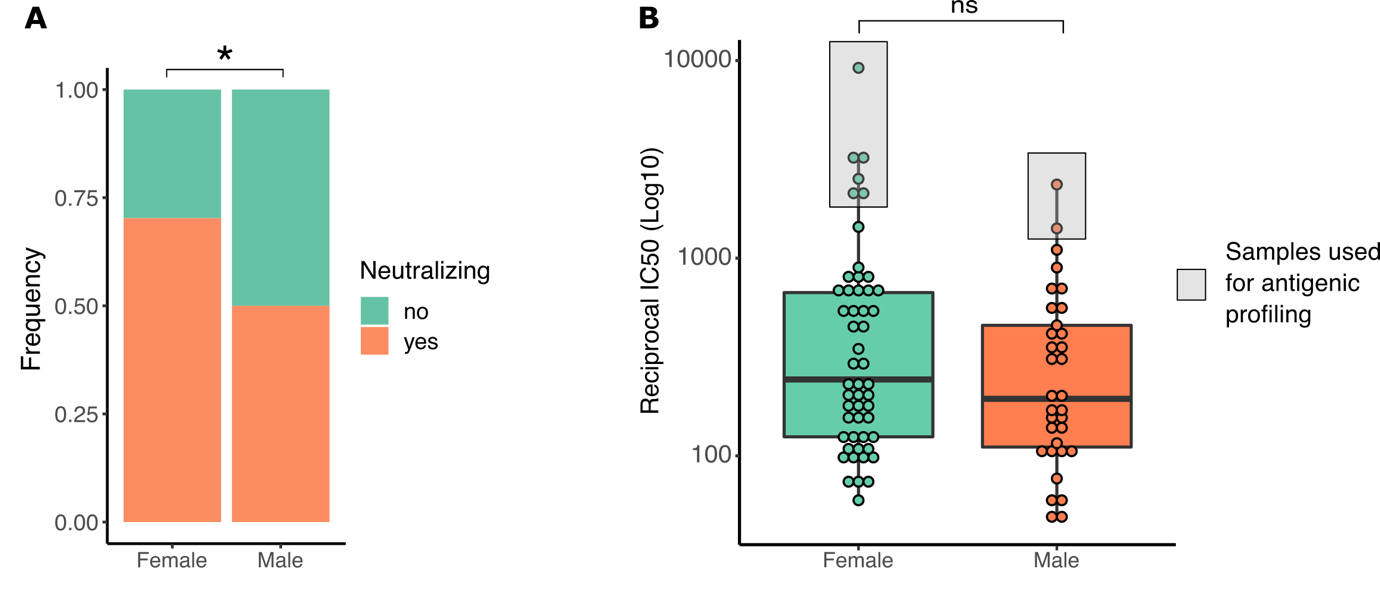

**Figure S2. Related to Figures 3 and 4. Screening for CVB3 neutralizing sera in healthy donors. (A)** Frequency of neutralizing sera, defined as having a reciprocal IC50 > 40, by sex. The prevalence of neutralizing samples was 60% (85/140). Female donors had higher seroprevalence (neutralizing samples female=52/74, male=33/66; p-value = 0.01605 by Fisher’s exact test). **(B)** Reciprocal IC50 values of neutralizing samples per sex (ns or p > 0.05 by Wilcoxon’s test). Samples used for the antigenic profiling experiments are highlighted with gray rectangles (top 6 female and top 2 male neutralizing sera).

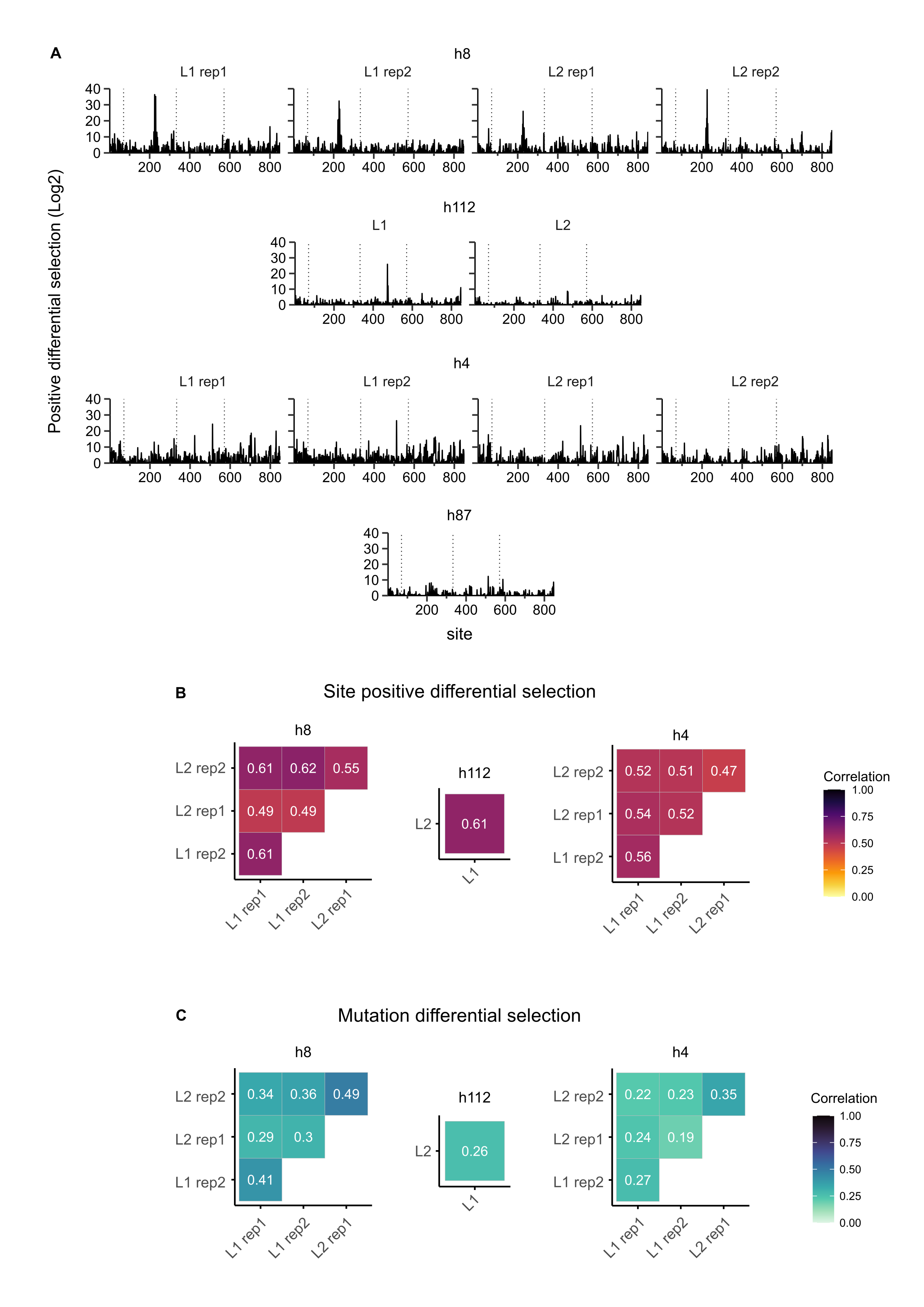

**Figure S3. Related to Figure 3. Technical and biological replicates of the mutational antigenic profiling of narrow human sera samples. (A)** Positive differential selection profiles of the narrow human sera samples (h8, h112, h4, h87). One (h112) or two (h4, h8) replicates (rep1 and rep2) were performed for two independent virus libraries: library 1 and library 2 (L1 and L2, respectively). Sample h87 was only analyzed with L2. **(B-C)** Matrix of Pearson correlation coefficients of site positive differential selection **(B)** and mutation differential selection **(C)** between biological and technical replicates.

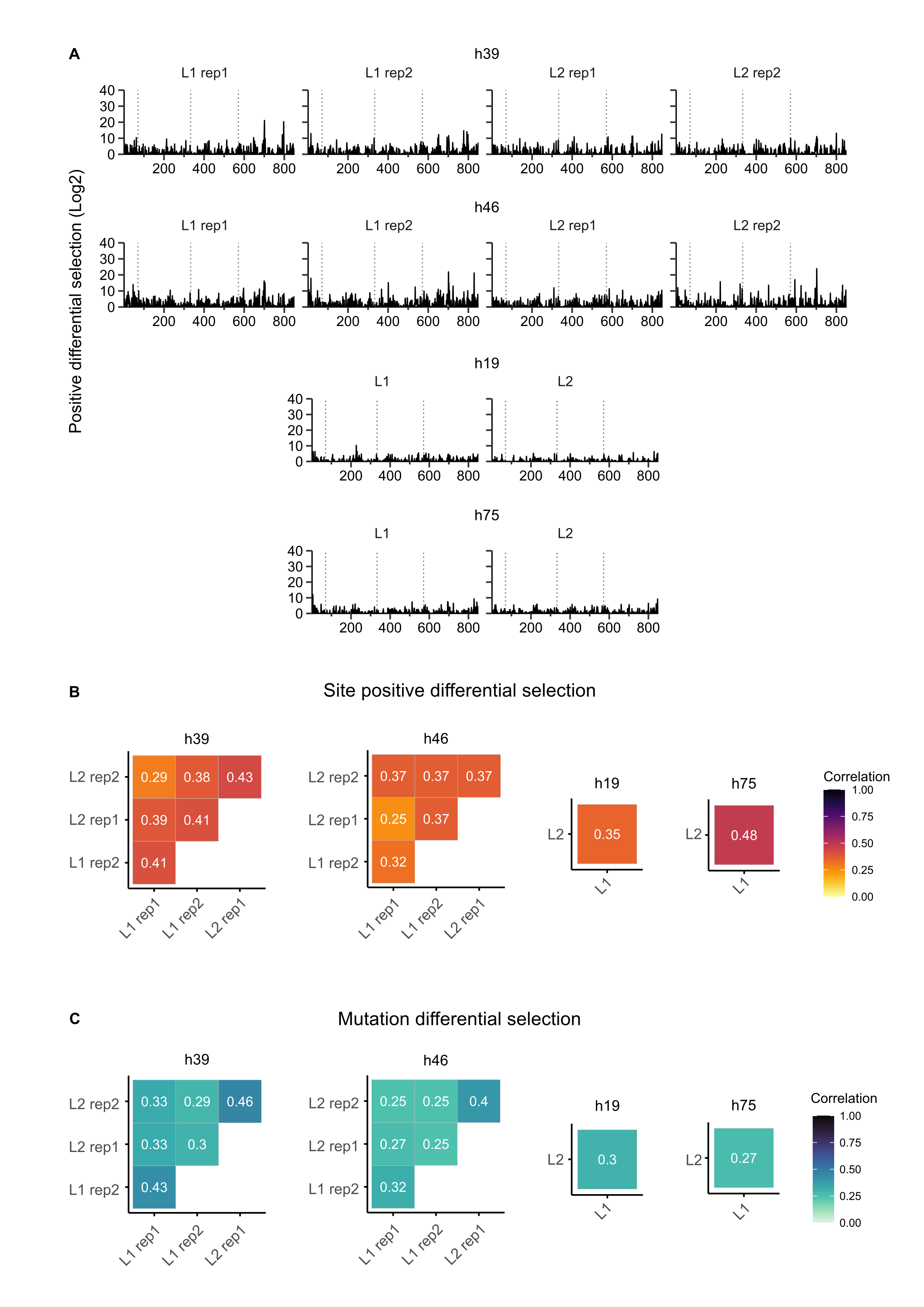

**Figure S4. Related to Figure 4. Technical and biological replicates of the mutational antigenic profiling of broad human sera. (A)** Positive differential selection profiles of broad human sera (h39, h46, h19, h75). One (h19, h75) or two (h39, h46) replicates (rep1 and rep2) were performed for two independent virus libraries: library 1 and library 2 (L1 and L2, respectively). **(B-C)** Matrix of Pearson correlation coefficients of site positive differential selection **(B)** and mutation differential selection **(C)** between biological and technical replicates.

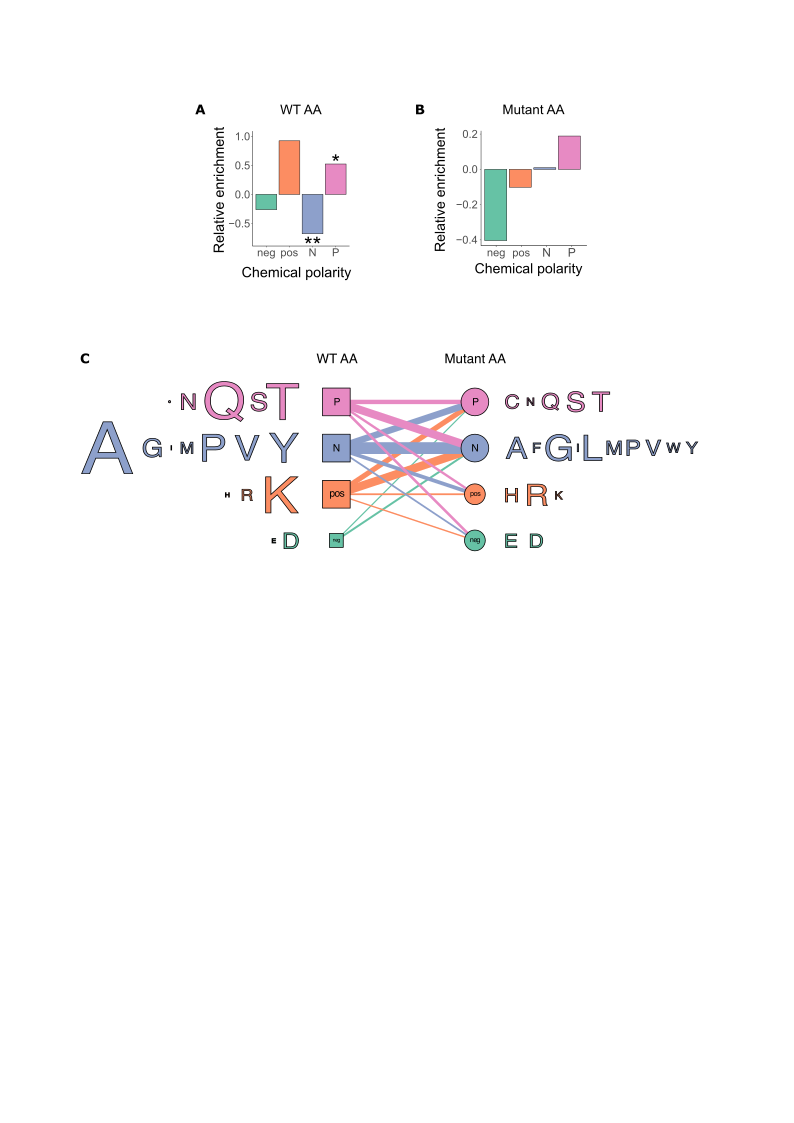

**Figure S5. Related to Figure 5. Analysis of mutations that confer escape from human polyclonal sera. (A)** The relative enrichment of the chemical polarity of amino acids in sites conferring escape versus surface exposed residues where no escape is observed. **(B)** The relative enrichment of the chemical polarity of individual amino acids conferring escape versus mutations in the same residues which do not confer escape. **(C)** Bipartite network analysis of WT and mutant AAs where escape is observed. Edge width is proportional to the number of occurrences. Vertex size is proportional to the number of connected edges. The size of amino acid letters is proportional to the number of occurrences. Color code according to physicochemical properties is the same as that used in Figure 5. P=polar, N=non-polar, pos=positively charged, neg=negatively charged.

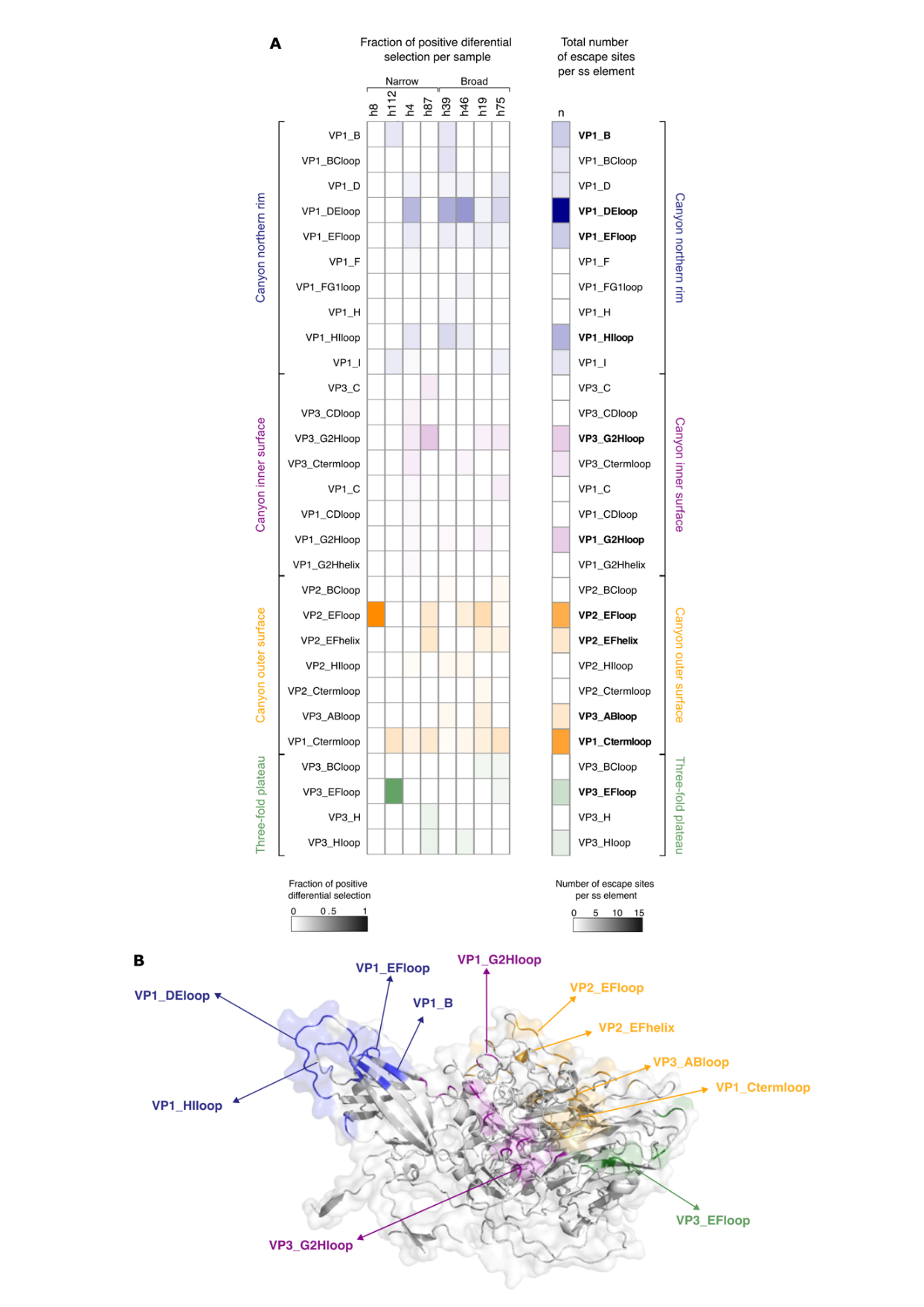

**Figure S6. Related to Figure 6. Structural distribution of top human escape sites. (A)** Heatmap representation of the total (right) or per sample (left) positive differential selection by secondary structure element and capsid region. Sites of escape were defined as surface exposed residues with a positive differential selection value above the mean+2SD value per sample (n=76, [https://‌github.com/‌RGellerLab/‌CVB3-Antigenic-Profiling/‌tree/‌main/‌10_‌analysis_‌sites_‌muts](https://github.com/RGellerLab/CVB3-Antigenic-Profiling/tree/main/10_analysis_sites_muts)). Secondary structure elements with a higher number of escape sites (n ≥ 3) are highlighted in bold. **(B)** Structural representation of the most prevalent secondary structure elements. The capsid monomer of the CVB3 strain Nancy is represented with the most prevalent secondary structure elements colored according to their structural location colored as in (A).

**Supplementary tables**

**Table S1. Related to Figures 1-4.** Surviving fraction of viral populations after neutralization with the mAb or the mouse and human polyclonal sera. Antibody concentration used and the number of replicates (rep) are indicated.

| **Library** | **Ab** | **rep** | **Ab concentration**  **(µL diluted in 100µL)** | **Surviving fraction** |
| --- | --- | --- | --- | --- |
| L1 | h4 | 1 | 1.25 | 2.50 |
| L1 | h4 | 2 | 1.25 | 2.97 |
| L1 | h8 | 1 | 1.25 | 0.73 |
| L1 | h8 | 2 | 1.25 | 0.91 |
| L1 | h39 | 1 | 5 | 1.70 |
| L1 | h39 | 2 | 5 | 3.40 |
| L1 | h46 | 1 | 5 | 0.87 |
| L1 | h46 | 2 | 5 | 1.11 |
| L1 | m1 | 1 | 5 | 4.09 |
| L1 | m1 | 2 | 5 | 11.69 |
| L1 | m2 | 1 | 5 | 43.74 |
| L1 | m2 | 2 | 5 | 12.18 |
| L1 | m3 | 1 | 5 | 10.53 |
| L1 | m3 | 2 | 5 | 12.45 |
| L1 | mAb | 1 | 7.5 | 3.91 |
| L1 | mAb | 2 | 7.5 | 1.62 |
| L1 | h112 | 1 | 5 | 0.92 |
| L1 | h19 | 1 | 5 | 0.44 |
| L1 | h75 | 1 | 5 | 0.59 |
| L2 | h4 | 1 | 1.25 | 1.81 |
| L2 | h4 | 2 | 1.25 | 2.69 |
| L2 | h8 | 1 | 1.25 | 2.84 |
| L2 | h8 | 2 | 1.25 | 1.22 |
| L2 | h39 | 1 | 5 | 4.12 |
| L2 | h39 | 2 | 5 | 4.77 |
| L2 | h46 | 1 | 5 | 2.43 |
| L2 | h46 | 2 | 5 | 1.75 |
| L2 | m1 | 1 | 5 | 3.31 |
| L2 | m1 | 2 | 5 | 2.39 |
| L2 | m2 | 1 | 5 | 8.67 |
| L2 | m2 | 2 | 5 | 12.01 |
| L2 | m3 | 1 | 5 | 5.56 |
| L2 | m3 | 2 | 5 | 5.69 |
| L2 | mAb | 1 | 7.5 | 2.32 |
| L2 | mAb | 2 | 7.5 | 1.75 |
| L2 | h87 | 1 | 1.5 | 0.31 |
| L2 | h112 | 1 | 5 | 2.37 |
| L2 | h19 | 1 | 5 | 1.14 |
| L2 | h75 | 1 | 5 | 2.75 |

**Table S2. Related to Figures 1 and 2. Main sites of escape of the mAb and mice antigenic profiles.** Surface exposed residues with a positive differential selection value above the mean+2SD per sample (SS: secondary structure element).

| **Sample(s)** | **Site** | **VP** | **Position** | **SS** |
| --- | --- | --- | --- | --- |
| m3 | 211 | 2 | 142 | VP2 EF loop |
| m2 | 212 | 2 | 143 | VP2 EF helix |
| m3 | 225 | 2 | 156 | VP2 EF loop |
| mAb, m3 | 227 | 2 | 158 | VP2 EF loop |
| mAb | 228 | 2 | 159 | VP2 EF loop |
| mAb | 229 | 2 | 160 | VP2 EF loop |
| mAb | 230 | 2 | 161 | VP2 EF loop |
| m3 | 232 | 2 | 163 | VP2 EF loop |
| m3 | 236 | 2 | 167 | VP2 EF loop |
| mAb | 242 | 2 | 173 | VP2 EF helix |
| m3 | 299 | 2 | 230 | VP2 H |
| m3 | 390 | 3 | 58 | VP3 AB loop |
| mAb | 391 | 3 | 59 | VP3 AB helix |
| mAb | 392 | 3 | 60 | VP3 AB helix |
| mAb, m3 | 393 | 3 | 61 | VP3 AB helix |
| mAb | 396 | 3 | 64 | VP3 AB loop |
| m3 | 401 | 3 | 69 | VP3 AB loop |
| m2, m3 | 513 | 3 | 181 | VP3 G2H loop |
| m3 | 541 | 3 | 209 | VP3 I |
| m1, m2, m3 | 650 | 1 | 80 | VP1 B |
| m1, m2, m3 | 652 | 1 | 82 | VP1 BC loop |
| m1, m2, m3 | 654 | 1 | 84 | VP1 BC loop |
| m1, m2, m3 | 655 | 1 | 85 | VP1 BC loop |
| m1 | 656 | 1 | 86 | VP1 BC loop |
| m2 | 698 | 1 | 128 | VP1 DE loop |
| m3 | 699 | 1 | 129 | VP1 DE loop |
| m2 | 701 | 1 | 131 | VP1 DE loop |
| m2 | 702 | 1 | 132 | VP1 DE loop |
| m1, m2, m3 | 723 | 1 | 153 | VP1 EF loop |
| m3 | 777 | 1 | 207 | VP1 G2H helix |
| m3 | 797 | 1 | 227 | VP1 HI loop |
| m1, m2, m3 | 800 | 1 | 230 | VP1 I |
| m2 | 830 | 1 | 260 | VP1 C-term loop |
| m1, m3 | 834 | 1 | 264 | VP1 C-term loop |
| m2, m3 | 840 | 1 | 270 | VP1 C-term loop |
| m3 | 843 | 1 | 273 | VP1 C-term loop |
| m3 | 846 | 1 | 276 | VP1 C-term loop |
| m2, m3 | 847 | 1 | 277 | VP1 C-term loop |
| m2 | 849 | 1 | 279 | VP1 C-term loop |

**Table S3. Related to Figures 3 and 4. Main sites of escape of the human antigenic profiles.** Surface exposed residues with a positive differential selection value above the mean+2SD per sample (SS: secondary structure element).

| **Samples** | **Site** | **VP** | **Position** | **SS** |
| --- | --- | --- | --- | --- |
| h39, h75 | 143 | 2 | 74 | VP2 BC loop |
| h46 | 208 | 2 | 139 | VP2 EF loop |
| h19 | 212 | 2 | 143 | VP2 EF helix |
| h75, h87 | 213 | 2 | 144 | VP2 EF helix |
| h8, h46, h75 | 221 | 2 | 152 | VP2 EF loop |
| h87 | 222 | 2 | 153 | VP2 EF loop |
| h8, h19 | 225 | 2 | 156 | VP2 EF loop |
| h8, h19 | 227 | 2 | 158 | VP2 EF loop |
| h8, h87 | 230 | 2 | 161 | VP2 EF loop |
| h19 | 231 | 2 | 162 | VP2 EF loop |
| h8 | 239 | 2 | 170 | VP2 EF loop |
| h19 | 242 | 2 | 173 | VP2 EF helix |
| h4, h39, h46 | 309 | 2 | 240 | VP2 HI loop |
| h19 | 331 | 2 | 262 | VP2 C-term loop |
| h19 | 388 | 3 | 56 | VP3 AB loop |
| h39 | 390 | 3 | 58 | VP3 AB loop |
| h19 | 401 | 3 | 69 | VP3 AB loop |
| h19, h75 | 410 | 3 | 78 | VP3 BC loop |
| h87 | 418 | 3 | 86 | VP3 C |
| h4 | 423 | 3 | 91 | VP3 CD loop |
| h112 | 473 | 3 | 141 | VP3 EF loop |
| h75, h112 | 475 | 3 | 143 | VP3 EF loop |
| h112 | 476 | 3 | 144 | VP3 EF loop |
| h4, h19, h75 | 512 | 3 | 180 | VP3 G2H loop |
| h87 | 513 | 3 | 181 | VP3 G2H loop |
| h87 | 515 | 3 | 183 | VP3 G2H loop |
| h87 | 524 | 3 | 192 | VP3 H |
| h46 | 535 | 3 | 203 | VP3 H Ioop |
| h87 | 536 | 3 | 204 | VP3 H Ioop |
| h46 | 561 | 3 | 229 | VP3 C-term loop |
| h4 | 564 | 3 | 232 | VP3 C-term loop |
| h39 | 647 | 1 | 77 | VP1 B |
| h39 | 648 | 1 | 78 | VP1 B |
| h39, h112 | 650 | 1 | 80 | VP1 B |
| h39 | 652 | 1 | 82 | VP1 BC loop |
| h39 | 656 | 1 | 86 | VP1 BC loop |
| h4, h75 | 661 | 1 | 91 | VP1 C |
| h4 | 666 | 1 | 96 | VP1 CD loop |
| h4 | 693 | 1 | 123 | VP1 D |
| h4, h39,  h46, h75 | 694 | 1 | 124 | VP1 D |
| h39 | 695 | 1 | 125 | VP1 DE loop |
| h4, h39, h75 | 696 | 1 | 126 | VP1 DE loop |
| h75 | 698 | 1 | 128 | VP1 DE loop |
| h4, h46 | 699 | 1 | 129 | VP1 DE loop |
| h39, h46 | 700 | 1 | 130 | VP1 DE loop |
| h4, h39, h46 | 701 | 1 | 131 | VP1 DE loop |
| h4, h39,  h46, h75 | 702 | 1 | 132 | VP1 DE loop |
| h4, h39, h46 | 703 | 1 | 133 | VP1 DE loop |
| h4, h46 | 704 | 1 | 134 | VP1 DE loop |
| h4, h39 | 705 | 1 | 135 | VP1 DE loop |
| h4, h19 | 706 | 1 | 136 | VP1 DE loop |
| h39 | 722 | 1 | 152 | VP1 EF loop |
| h4, h39,  h19, h75 | 723 | 1 | 153 | VP1 EF loop |
| h4, h46 | 724 | 1 | 154 | VP1 EF loop |
| h4 | 740 | 1 | 170 | VP1 F |
| h46 | 742 | 1 | 172 | VP1 FG1 loop |
| h19 | 772 | 1 | 202 | VP1 G2H loop |
| h39 | 774 | 1 | 204 | VP1 G2H loop |
| h4 | 779 | 1 | 209 | VP1 G2H helix |
| h4 | 782 | 1 | 212 | VP1 G2H loop |
| h39 | 790 | 1 | 220 | VP1 H |
| h39 | 793 | 1 | 223 | VP1 HI loop |
| h4 | 796 | 1 | 226 | VP1 HI loop |
| h4, h39, h46 | 797 | 1 | 227 | VP1 HI loop |
| h4, h39, h46 | 798 | 1 | 228 | VP1 HI loop |
| h75, h112 | 800 | 1 | 230 | VP1 I |
| h4 | 802 | 1 | 232 | VP1 I |
| h4, h75 | 827 | 1 | 257 | VP1 C-term loop |
| h4, h46, h75 | 829 | 1 | 259 | VP1 C-term loop |
| h46 | 831 | 1 | 261 | VP1 C-term loop |
| h39, h75 | 840 | 1 | 270 | VP1 C-term loop |
| h75, h87 | 843 | 1 | 273 | VP1 C-term loop |
| h19 | 846 | 1 | 276 | VP1 C-term loop |
| h87, h112 | 847 | 1 | 277 | VP1 C-term loop |
| h112 | 848 | 1 | 278 | VP1 C-term loop |
| h19 | 849 | 1 | 279 | VP1 C-term loop |

**Table S4. Related to Figure 5. Comparison between different machine learning algorithms.**

|  |  |  | **Prior Prob.**  **+**  **Prob. Recal** | **Prior Prob.**  **+**  **No Prob. Recal** | **No Prior Prob.**  **+**  **Prob. Recal** | **No Prior Prob.**  **+**  **No Prob. Recal** |
| --- | --- | --- | --- | --- | --- | --- |
| **Machine**  **Learning**  **Algorithm** | **Neural**  **Network** | **Accuracy** | $95.2\pm0.6\%$ | $95.1\pm0.6\%$ | $91.8\pm0.7\%$ | $92.9\pm0.7\%$ |
|  |  | **F_1_-score** | **no escape:**  $97.5\pm0.3\%$  **escape**:  $17\pm5\%$ | **no escape:**  $97.5\pm0.3\%$  **escape**:  $12\pm4\%$ | **no escape**:  $95.7\pm0.5\%$  **escape**:  $18\pm3\%$ | **no escape**:  $96.3\pm0.4\%$  **escape**:  $22\pm4\%$ |
|  |  | **Precision** | **no escape**:  $96.7\pm0.5\%$  **escape**:  $24\pm8\%$ | **no escape**:  $96.5\pm0.5\%$  **escape**:  $19\pm8\%$ | **no escape**:  $97.0\pm0.5\%$  **escape**:  $14\pm4\%$ | **no escape**:  $97.1\pm0.5\%$  **escape**:  $18\pm4\%$ |
|  |  | **ROC AUC** | **no escape**:  $81\pm1\%$  **escape**:  $81\pm1\%$ | **no escape**:  $83\pm1\%$  **escape**:  $83\pm1\%$ | **no escape**:  $82\pm1\%$  **escape**:  $82\pm1\%$ | **no escape**:  $81\pm1\%$  **escape**:  $81\pm1\%$ |
|  | **Support**  **Vector**  **Machine** | **Accuracy** | $95.4\pm0.5\%$ | $95.2\pm0.6\%$ | $90.1\pm0.8\%$ | $92.3\pm0.7\%$ |
|  |  | **F_1_-score** | **no escape**:  $97.6\pm0.3\%$  **escape**:  $28\pm5\%$ | **no escape**:  $97.5\pm0.3\%$  **escape**:  $29\pm5\%$ | **no escape**:  $95.0\pm0.4\%$  **escape**:  $26\pm3\%$ | **no escape**:  $95.9\pm0.4\%$  **escape**:  $29\pm4\%$ |
|  |  | **Precision** | **no escape**:  $97.1\pm0.4\%$  **escape**:  $34\pm8\%$ | **no escape**:  $97.1\pm0.4\%$  **escape**:  $32\pm7\%$ | **no escape**:  $97.7\pm0.4\%$  **escape**:  $19\pm3\%$ | **no escape**:  $97.7\pm0.4\%$  **escape**:  $22\pm4\%$ |
|  |  | **ROC AUC** | **no escape**:  $84.8\pm0.9\%$  **escape**:  $84.8\pm0.9\%$ | **no escape**:  $85.1\pm0.9\%$  **escape**:  $85.1\pm0.9\%$ | **no escape**:  $85.5\pm0.9\%$  **escape**:  $85.5\pm0.9\%$ | **no escape**:  $84.8\pm0.9\%$  **escape**:  $84.8\pm0.9\%$ |
|  | **Random**  **Forest** | **Accuracy** | $95.7\pm0.5\%$ | $96.3\pm0.5\%$ | $92.8\pm0.7\%$ | $93.2\pm0.7\%$ |
|  |  | **F_1_-score** | **no escape**:  $97.8\pm0.3\%$  **escape**:  $26\pm5\%$ | **no escape**:  $98.1\pm0.3\%$  **escape**:  $0\%$ | **no escape**:  $96.2\pm0.4\%$  **escape**:  $36\pm4\%$ | **no escape**:  $96.4\pm0.4\%$  **escape**:  $36\pm4\%$ |
|  |  | **Precision** | **no escape**:  $97.0\pm0.5\%$  **escape**:  $37\pm9\%$ | **no escape**:  $96.3\pm0.5\%$  **escape**:  $0\%$ | **no escape**:  $98.1\pm0.4\%$  **escape**:  $27\pm4\%$ | **no escape**:  $98.0\pm0.4\%$  **escape**:  $28\pm5\%$ |
|  |  | **ROC AUC** | **no escape**:  $90.7\pm0.8\%$  **escape**:  $90.6\pm0.8\%$ | **no escape**:  $90.1\pm0.8\%$  **escape**:  $89.9\pm0.8\%$ | **no escape**:  $90.7\pm0.8\%$  **escape**:  $90.6\pm0.8\%$ | **no escape**:  $90.1\pm0.8\%$  **escape**:  $89.9\pm0.8\%$ |

**Table S5. Related to Figure 5. Classification metrics of the Random Forest classifiers tested.**

|  | **All features +**  **no reclassification** | | **All features +**  **reclassification (27 cases)** | | **Top 7 features +**  **reclassification** | |
| --- | --- | --- | --- | --- | --- | --- |
|  | **Training** | **Testing** | **Training** | **Testing** | **Training** | **Testing** |
| **Accuracy** | $99.72\pm0.06\%$ | $95.7\pm0.5\%$ | $99.84\pm0.04\%$ | $96.1\pm0.5\%$ | $99.72\pm0.06\%$ | $96.8\pm0.5\%$ |
| **F_1_-score** | **no escape:**  $99.72\pm0.06\%$  **escape:**  $99.72\pm0.06\%$ | **no escape:**  $97.8\pm0.3\%$  **escape:**  $26\pm5\%$ | **no escape:**  $99.84\pm0.04\%$  **escape:**  $99.84\pm0.04\%$ | **no** **escape:**  $98.0\pm0.3\%$  **escape:**  $50\pm5\%$ | **no** **escape:**  $99.72\pm0.06\%$  **escape:**  $99.72\pm0.06\%$ | **no** **escape:**  $98.4\pm0.2\%$  **escape:**  $50\pm5\%$ |
| **Precision** | **no escape:**  $100\%$  **escape:**  $99.5\pm0.1\%$ | **no** **escape:**  $97.07\pm0.4\%$  **escape:**  $37\pm9\%$ | **no** **escape:**  $100\%$  **escape:**  $99.69\pm0.09\%$ | **no** **escape:**  $97.7\pm0.4\%$  **escape:**  $54\pm7\%$ | **no** **escape:**  $100\%$  **escape:**  $99.5\pm0.01\%$ | **no** **escape:**  $97.4\pm0.4\%$  **escape:**  $72\pm8\%$ |
| **ROC AUC** | **no** **escape:**  $99.99\pm0.01\%$  **escape:**  $99.99\pm0.01\%$ | **no** **escape:**  $90.7\pm0.8\%$  **escape:**  $90.6\pm0.8\%$ | **no** **escape:**  $99.999\pm0.002\%$  **escape:**  $99.999\pm0.002\%$ | **no** **escape:**  $93.4\pm0.7\%$  **escape:**  $93.3\pm0.7\%$ | **no** **escape:**  $99.993\pm0.009\%$  **escape:**  $99.99\pm0.01\%$ | **no** **escape:**  $93.2\pm0.7\%$  **escape:**  $93.1\pm0.7\%$ |
